# Image Analysis Techniques for In Vivo Quantification of Cerebrospinal Fluid Flow

**DOI:** 10.1101/2023.07.20.549937

**Authors:** Daehyun Kim, Yiming Gan, Maiken Nedergaard, Douglas H. Kelley, Jeffrey Tithof

## Abstract

Over the last decade, there has been a tremendously increased interest in understanding the neurophysiology of cerebrospinal fluid (CSF) flow, which plays a crucial role in clearing metabolic waste from the brain. This growing interest was largely initiated by two significant discoveries: the glymphatic system (a pathway for solute exchange between interstitial fluid deep within the brain and the CSF surrounding the brain) and meningeal lymphatic vessels (lymphatic vessels in the layer of tissue surrounding the brain that drain CSF). These two CSF systems work in unison, and their disruption has been implicated in several neurological disorders including Alzheimer’s disease, stoke, and traumatic brain injury. Here, we present experimental techniques for *in vivo* quantification of CSF flow via direct imaging of fluorescent microspheres injected into the CSF. We discuss detailed image processing methods, including registration and masking of stagnant particles, to improve the quality of measurements. We provide guidance for quantifying CSF flow through particle tracking and offer tips for optimizing the process. Additionally, we describe techniques for measuring changes in arterial diameter, which is an hypothesized CSF pumping mechanism. Finally, we outline how these same techniques can be applied to cervical lymphatic vessels, which collect fluid downstream from meningeal lymphatic vessels. We anticipate that these fluid mechanical techniques will prove valuable for future quantitative studies aimed at understanding mechanisms of CSF transport and disruption, as well as for other complex biophysical systems.

## 1 Introduction

Cerebrospinal fluid (CSF) envelops the brain and spinal cord, acting as a cushion that provides buoyancy [1] and supplying nutrients [1, 2]. Growing evidence suggests that CSF also plays an important role in metabolic waste removal [2–5]. Since the brain parenchyma is devoid of lymphatic vessels, researchers have speculated for decades that CSF circulation may serve a pseudo-lymphatic role [6]. Tracer injection experiments from past decades [7–9] demonstrated that solute exchange between the CSF and interstitial fluid (ISF) occurs at rates faster than diffusion alone, indicating the presence of bulk flow through perivascular spaces (PVSs), which are annular channels surrounding vasculature throughout the brain. Several high-profile discoveries from the past decade [4, 10–16] have inspired growing interest in CSF research due to their profound implications for improving health. However, many questions remain regarding the direction, key compartments, and mechanisms of CSF flow throughout the central nervous system.

The precise details of solute exchange between CSF and ISF are not well-established, with multiple competing hypotheses in the literature. The intramural periarterial drainage (IPAD) hypothesis posits that basement membranes (i.e., vessel walls) of cerebral capillaries and arteries are the main efflux route by which ISF exits the brain [17–19]. This hypothesis is largely motivated by observations of amyloid-*β* (a protein waste molecule linked to neurodegenerative diseases) aggregation on arteries, which is a hallmark feature of cerebral amyloid angiopathy [20]. Carare et al [21] reported that soluble tracers diffuse through the brain parenchyma and then drain through basement membranes, indicating ISF is transported retrograde to blood flow. However, it is important to note that the vast majority of experimental evidence supporting the IPAD hypothesis is derived from ex vivo analysis of fixed tissue, which is subject to anatomical changes and irregular flows that occur during the fixation process [12]. Alternatively, the glymphatic (glial-lymphatic) hypothesis suggests that CSF follows a pathway that is anterograde to blood flow, flowing inward along PVSs of arteries, continuing through the brain interstitium, then exiting via venous PVSs [4, 22, 23]. Within the framework of either hypothesis, arterial pulsations are widely acknowledged as a likely mechanism driving bulk CSF/ISF flow [4, 12, 24, 25]. Several prior studies have analytically and numerically modeled arterial pulsations to investigate the feasibility of this driving mechanism [26–29], which has led to a third hypothesis: that arterial pulsations generate oscillatory (net zero) flow that enhances transport via Taylor dispersion [28]. We point the reader to several review articles for a more extensive discussion of CSF/ISF transport hypotheses [30–35].

CSF drains from the skull through multiple parallel efflux routes including along cranial nerves that penetrate the cribriform plate [36–39], through basal and dorsal meningeal lymphatic vessels [40, 41], to the spinal canal [42], and directly into the blood at the superior sagittal sinus [43]; however, the exact contribution of each route is not well understood [44]. Further complicating the matter, recent research suggests that the rate and distribution of CSF production and outflow varies depending on the circadian rhythm [45, 46]. Regardless of the exact details of outflow dynamics, it has long been appreciated that a substantial portion of CSF drainage eventually reaches cervical lymphatic vessels located in the neck [47, 48], and the discovery of meningeal lymphatic vessels supports the idea that the lymphatic system constitutes a significant CSF outflow route. Ultimately, many open questions remain due to experimental challenges and technical limitations associated with probing small length scales deep inside or adjacent to the skull.

Numerous experimental approaches exist for measuring CSF flow [35], but each has different benefits and drawbacks. In this article, we describe image analysis techniques for quantifying CSF flow, applied to time series of two-dimensional (2D) images that were recorded using two-photon microscopy (TPM). TPM offers excellent spatial and temporal resolution (as high as about 0.5 µm/pixel at 60 Hz), enabling detailed fluid mechanical analysis of CSF dynamics through PVSs at the surface of the brain [12, 15, 49, 50] or through cervical lymphatic vessels in the neck [50, 51]. An important appealing feature of TPM is that it relies on *two-photon* fluorophore excitation which depends quadratically on the incident photon flux, and thus by focusing a laser at a precise location, fluorescence is achieved in a tiny volume of approximately 1 µm^3^ [52]. By rastering pixel-by-pixel throughout a plane, 2D images are achieved at a precise depth with negligible fluorophore excition above/below that plane; this is in contrast to other microscopy techniques (e.g., confocal). In 2018, Mestre et al [12] leveraged TPM to demonstrate that CSF flows parallel (not anti-parallel) to blood through approximately 40 µm wide PVSs at the surface of the brain, and the flow pulses in synchrony with the cardiac cycle [12] (anesthetized mice have a heart rate on the order of 5 Hz, so a Nyquist rate < 10 Hz is needed to resolve the flow pulsatility). Subsequently, these techniques were further refined and utilized to quantify changes in CSF dynamics following stroke [15], cardiac arrest [53], and traumatic brain injury [50]. We posit that as research interest in CSF flow continues to grow due to its increasingly apparent relevance to a variety of chronic and acute neurological conditions [54, 55], *in vivo* optical measurement techniques in animal models will continue to provide valuable insight into CSF dynamics. However, it is important to note the limitations of TPM: this technique requires invasive surgery to place a cranial window enabling optical access to the brain, the field of view is limited, and imaging is restricted to the top few hundred µm of the cortex since brain tissue scatters light. Insights from high-resolution measurements, obtained using techniques such as those described in this article, will likely prove most valuable when integrated into a comprehensive framework that includes other modalities for measuring CSF transport, such as magnetic resonance imaging (MRI) [56–59], transcranial brain-wide epifluorescence microscopy [60, 61], and *ex vivo* tissue fixation with histological staining [7–9, 17–19, 21]. For a more comprehensive discussion of various measurement techniques, we point the reader to Bohr et al [35].

In this article, we provide an overview of image analysis techniques that enable quantitative characterization of CSF dynamics. Our MATLAB-based scripts, along with a short time series of images and a concise working example, are freely available online [62]. A key component of the analysis involves particle tracking velocimetry (PTV), which has been used extensively for quantifying biological phenomena including respiratory flows [63–65], cell migration [66–68], and blood flow [69–71]. We do not provide any discussion of surgical techniques required prior to imaging CSF flow *in vivo*; rather, we direct the reader to references [61, 72].

This article is structured as follows. First, section 2 provides a brief overview of experimental methods and image analysis techniques described in this article. We then discuss image preprocessing in section 3, including image registration and masking stagnant microspheres, which both improve the reliability of CSF flow measurements. We provide insight and recommendations for choosing near-optimal tracking parameters in section 4. Then we introduce techniques for quantifying changes in vessel diameter, which are thought to contribute to CSF transport, in section 5. Next, we demonstrate in section 6 how the aforementioned techniques can be adapted and applied to measure fluid flow through cervical lymphatic vessels. Finally, a summary and conclusions are provided in section 7. An appendix is also included (section A) which presents an uncertainty analysis of PTV based on TPM imaging.

## 2 Overview of Experimental Methods

The image analysis techniques presented in this article require injection of fluorescent tracers to visualize different fluid compartments in the brain. To enable quantitative measurement of CSF flow via PTV, fluorescent microspheres (FluoSpheres, diameter 1.0 µm) were injected into the CSF, which is usually performed at the cisterna magna (an accessible region near the posterior skull base) [12, 72, 73]. Although some researchers have raised concerns that invasive tracer injections increase the intracranial pressure and may result in non-physiological flow [74–78], Raghunandan et al [49] demonstrated that flow velocities in PVSs are unchanged (compared to prior experimental protocols) when injections are performed using a dual syringe that injects and withdraws fluid, minimally disrupting the intracranial pressure. Fluorescent microspheres in the CSF appear as green dots in Fig. 1a. Experiments typically also include injection of fluorescent dye into the blood for visualization, which may be an intravenous or retro-orbital injection [79]. Visualizing the blood compartment is valuable for inferring the approximate location of the PVS (by definition, PVSs are adjacent to blood vessels and the vascular wall forms the inner boundary of each PVS). Such visualization also facilitates image registration and enables measurement of arterial pulsations, both described below. We note that here, as well as in prior publications [12, 15, 53], the color channels for CSF microspheres (green) and blood (red) have been swapped from their true fluorescence (red and green, respectively) to facilitate an intuitive representation of blood with the color red.

**Fig. 1.**
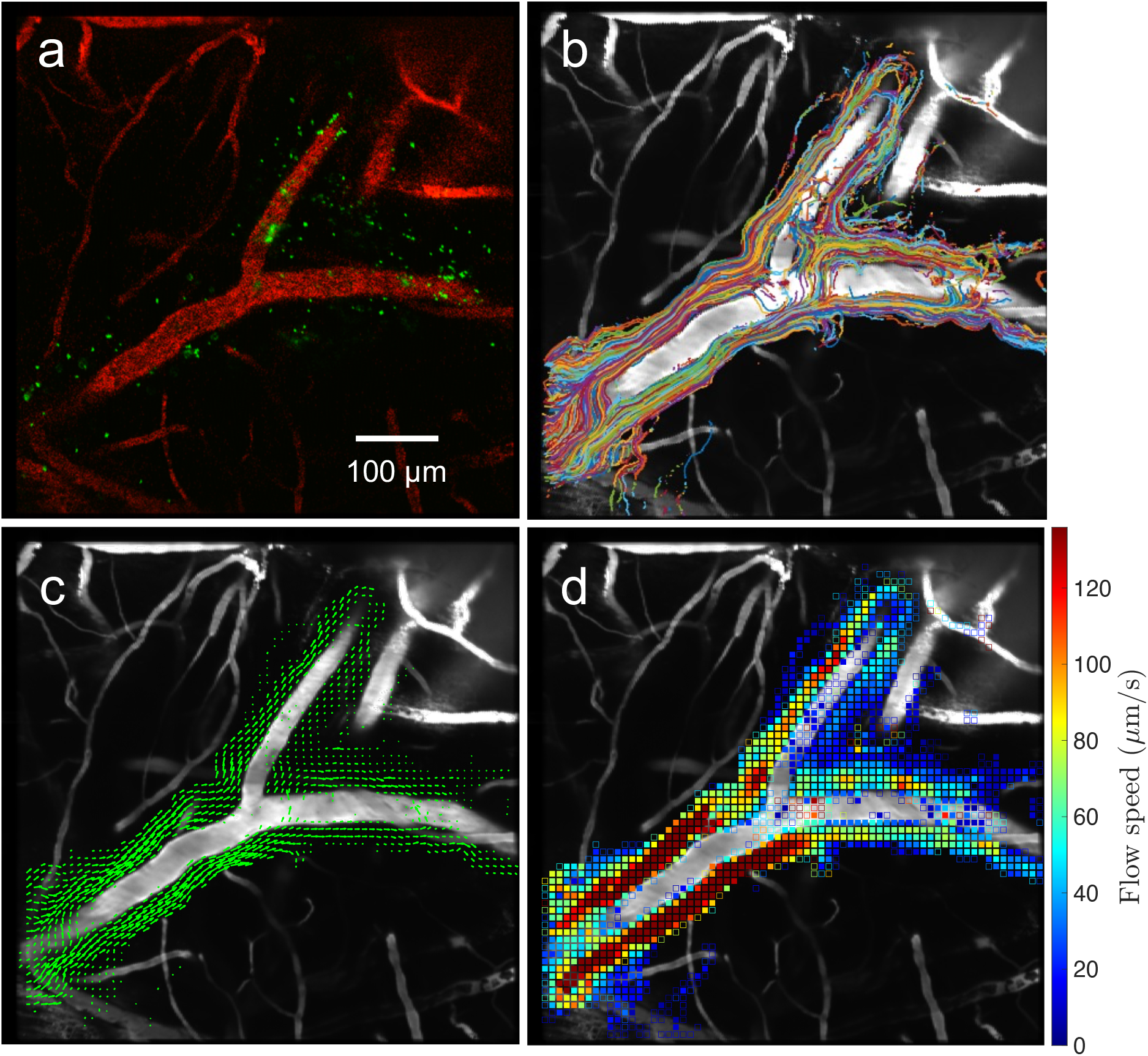
An example of image analysis to quantify CSF flow through PVSs at the surface of a mouse brain. Fluorescent tracers are injected into the blood (bovine serum albumin) and CSF (1 µm polystyrene microspheres) and then recorded *in vivo* using two-photon microscopy by imaging through a cranial window. **(a)** A snapshot of a pial artery (red) and fluorescent microspheres (green) acquired at 30 Hz. **(b)** Superimposed trajectories of the microspheres obtained from PTV, which can be used to visualize the size of the PVSs. **(c-d)** Time-averaged (c) velocity field and (d) flow speed quantifying the net CSF transport.

After a time series of microspheres flowing through the PVSs is recorded, we perform Lagrangian particle tracking using a predictive algorithm [80, 81]. For a given snapshot (e.g., Fig. 1a), particles are first detected based on a pixel intensity threshold and the centroid is identified for each contiguous region above that threshold. Identified particles are then linked in time by matching a given centroid to a nearby centroid in the next frame using kinematic predictions. The maximum search radius for linking sequential centroids in time is user-specified and should be chosen carefully (see below). Once particle tracking is completed, we plot the superimposed tracks from the entire time series recording, which provides a visual assessment of the PVS size and shape (Fig. 1b). PVS size/shape can also be characterized through infusion of fluorescent dyes (e.g., dextran, bovine serum albumin) into the CSF [4, 12, 82], but these small molecules can cross from the PVS into the brain interstitium, potentially obscuring interpretation of the location of the PVS outer boundary (which is formed by astrocyte endfeet). Finally, velocity measurements from the entire time series are spatially binned and averaged to obtain the mean CSF velocity (Fig. 1c) and the magnitude of each vector provides the flow speed (Fig. 1d). To improve the accuracy and reliability of the resulting velocity measurements, we perform image preprocessing, including image registration and masking of stagnant microspheres; these two procedures are presented in the next section.

## 3 Image Preprocessing

As described in the previous section, we obtain measurements of net CSF velocity in PVSs at the surface of the brain by performing *in vivo* PTV followed by spatial binning and averaging of all measurements.

Since these measurements are performed *in vivo*, some amount of relative motion between the microscope and the living mouse is practically unavoidable. Our recordings are necessarily obtained in the fixed reference frame of the microscope and hence should be transformed to a fixed reference frame of the region of the brain under consideration. We perform this transformation via image registration, which is described next in section 3.1. Afterward, in section 3.2, we discuss issues of microsphere aggregation and adherence to PVS boundaries, then present methods for masking these stagnant particles. Both image registration and masking techniques have been applied in prior studies [83–87] to improve accuracy of particle tracking and/or particle image velocimetry.

### 3.1 Registration

A significant challenge faced when performing *in vivo* CSF imaging is that the field of view may translate, which may occur in the *x−y* plane or perpendicular to it (along *z*), over short or long time scales. These shifts may be attributed to: (i) cardiac/respiratory cycles (short time scales), (ii) thermal expansion of imaging components (long), and/or (iii) small amounts of brain swelling resulting from cranial window placement (long). Five snapshots from an imaging time series are shown in Fig. 2a, for which subtle, long-time-scale translations occurred during the recording. This shift is perhaps most apparent from features in the top left corner, where some vessels are out of the field of view at *t* = 0, but become visible at *t* = 337*s*. This appearance of a new vascular structure near the boundary is indicative of a shift in the *x − y* plane. Additionally, there is an x-shaped configuration of vessels near the bottom left corner which is most apparent at times *t* = 337 and 1350 s. Appearance/disappearance of vasculature away from the boundaries is indicative of a shift in the imaging plane along the *z*-direction (recall that for TPM, images are acquired at a precise plane with depth of about 1 µm). There are limited means for accounting for a shift along *z*. Translations of the field of view, if not corrected for, will result in erroneous velocity measurements for two reasons: (i) motion of a particle between sequential frames will be due to both fluid flow and imaging plane translation, and (ii) PVSs will appear wider with blunted time-averaged peak flow speeds.

**Fig. 2.**
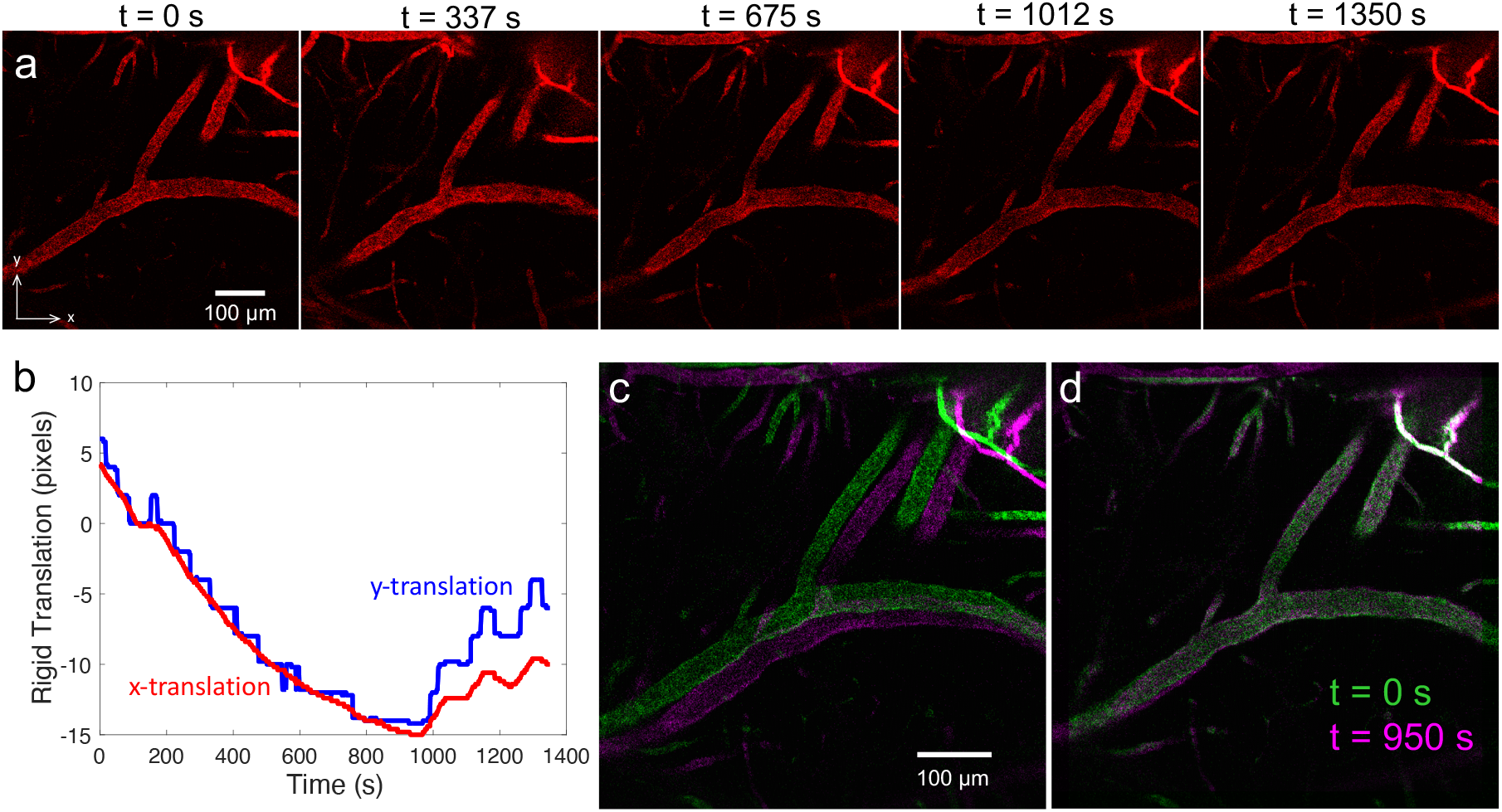
Example illustrating image registration in the vicinity of a pial artery. **(a)** Snapshots from a time series which show a small, gradual translation in the field of view. **(b)** Plot of the *x*-directional and *y*-directional translations that occur in time series, computed via cross-correlation with a reference frame; these translations are used for image registration. **(c-d)** Comparison of images at *t* = 0 s (green) and *t* = 950 s (magenta) (c) before and (d) after registration is performed.

To register images, we perform a spatial cross-correlation between a reference snapshot and every other snapshot in the time series to determine the transformation necessary to shift each image in the time series to a common reference frame. For flow through PVSs, we have found that registration that only accounts for rigid translation is adequate (i.e., rotation and deformation are unnecessary). We choose the reference snapshot typically as a frame (or time-average a short segment of frames to reduce noise) from a relatively stable instant in the recording, which we identify by visually inspecting a sped up animation of the TPM time series. After obtaining the cross-correlation peak for each frame, all frames in all channels are registered to the reference frame by applying a rigid translation (Fig. 2b). We note that it is necessary to increase the dimensions of the registered images by a number of pixels equal to the absolute value of the difference in maximum and minimum translations along each direction (e.g., based on Fig. 2b, approximately 4 *−* (*−*15) = 19 pixels must be added). This is so that the original image may be translated within the bounds of the registered image domain.

A successful application of image registration is illustrated in Fig. 2c-d wherein two snapshots of a pial artery are superimposed with different colors indicating time: green at *t* = 0 s and magenta at *t* = 950 s. Figure 2c shows these snapshots prior to registration, where there is a clear offset in the position of the vasculature; in Fig. 2d, which shows the post-registration snapshots, the colocalization of green and magenta indicates that the alignment is excellent. This figure also conveys details of the translation that occurred during imaging: near the top left of Fig. 2d, a segment of vasculature is purely magenta with no green, indicating that segment became visible at later times (*t* = 950 s) as the field of view translated in the positive *y*-direction.

### 3.2 Masking

As the fluorescent microspheres advect with the CSF flow, some of these particles adhere to the boundaries of the PVS. Once this occurs, they rarely become unstuck. In general, the reasons for particle aggregation and adherence to tissue are not well understood, but may be due to surface charge interactions or regions with fibrous obstructions forming a mesh that particles become trapped in (e.g., stomata that enable CSF exchange between pial PVSs and the subarachnoid space [88]). Figure 3a shows a snap-shot with an inset that provides an enlarged view of a region with a cluster of stuck particles (yellow box). Stagnant particles may interfere with measurement of CSF flow velocity by generating erroneous zero-velocity measurements. Hence, masking the stuck region will improve the measurement accuracy.

**Fig. 3.**
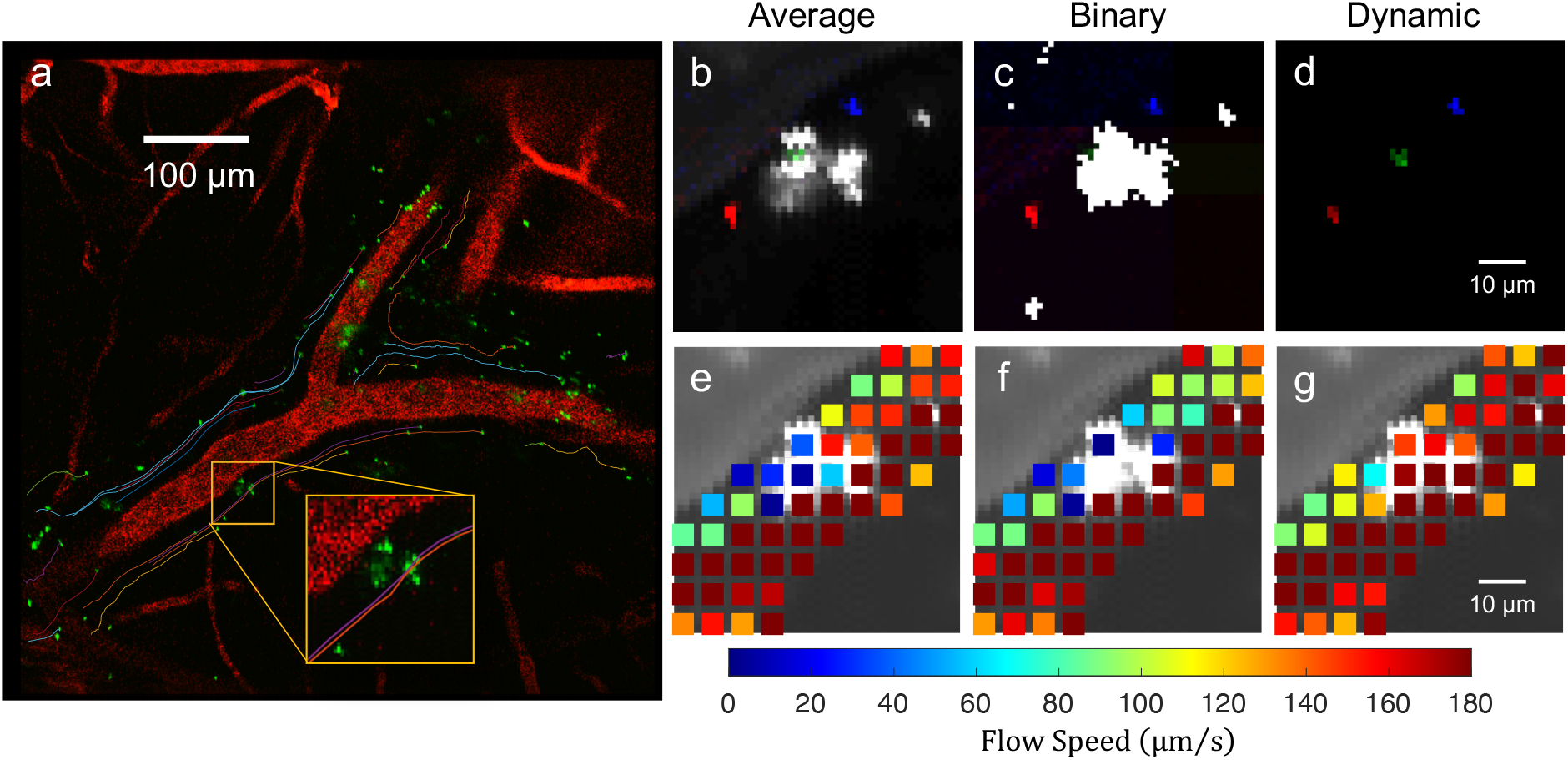
Comparison of masking methods. **(a)** Image of a pial artery with colored curves indicating the tracking of fluorescent microspheres flowing in the adjacent PVSs. No background has been subtracted for this image. (Inset) A zoomed-in image of a region containing a stagnant microsphere aggregate, which may lead to erroneous measurements of zero velocity if not properly masked. **(b-d)** Plots characterizing three different masking techniques (average, binary, and dynamic masking), with the mask superimposed in white/gray. Three sequential unmasked images are also superimposed in red, green, and blue indicating the trajectory of a flowing particle. **(e-g)** Velocity measurements obtained from particle tracking for a long time series with the indicated masking technique applied. (e) Average masking incompletely masks the region with stuck particles, leading to erroneous low-velocity measurements. (f) Binary masking completely removes measurements from the portion of the domain with stuck particles. (g) Dynamic masking leads to the best results with large velocities near the center of the channel, as expected, and no measurement voids, as in (f).

A variety of masking approaches have been implemented in velocity measurement applications [89]. One common masking approach is to create a “background image” by time-averaging an entire time series, then that background image is subtracted from all frames in the time series, enabling isolation of only the moving components. Figure 3b shows the time-averaged background image for the inset shown in Fig. 3a, where the pixel intensity of the background image varies from 0 to 255 (i.e., the grayscale background image has black, gray, and white values). The white regions of the image correspond to particles that stagnated early in the imaging times series, whereas gray regions correspond to regions where particle stagnation occurred later. For comparison, three sequential background-subtracted images are superimposed in red, green, and blue, indicating the time evolution of a free flowing particle that does not adhere to the stagnant cluster. This figure illustrates that masking with a standard time-average background image may or may not obscure a free-flowing particle. Detrimentally, late in the time series when the particle aggregate becomes large, the background image intensity may not be great enough to mask the entire stagnant aggregate, leading to zero velocity measurements. Such measurements are erroneous, as particles are still clearly flowing through the PVS at a slightly different depth from the stagnant aggregate.

A second masking approach is to replace the time-averaged background image with a binary background image. The purpose of generating a binary background image is to completely mask any region with stagnant particles. Figure 3c shows the background image obtained when a low pixel intensity threshold is used to binarize the background image presented above (i.e., values above or below the threshold become 255 or zero, respectively). Again, three sequential background-subtracted images are superimposed in red, green, and blue, but the green particle is not visible, indicating that this free flowing particle is indeed removed by this mask. However, at late times, this approach will satisfactorily mask regions with stagnant aggregates, avoiding erroneous zero velocity measurements. Note that this approach was used in Mestre et al [12] and explains why Figs. 1d and 5a in that manuscript have interspersed voids in the flow speed heatmap plots.

A third and final approach is dynamic masking whereby a separate background image is generated and subtracted from each image in the time series based on the average of the closest *n* frames in time, where *n* is user-specified (usually corresponding to about 5 s). In this approach, at any given frame, the spatial region that is masked will have a size and intensity that is much closer to optimal for masking the stagnant region. Hence, stagnant particles are masked and flowing particles are visible (Fig. 3d). This often even allows for tracking of particles that pass directly over/under a stagnant region, wherein the flowing particle transiently increases the local image intensity. The superiority of dynamic background masking is readily apparent when comparing Fig. 3e-g, where average masking yields reduced flow speeds near the center of the channel (Fig. 3e), binary masking yields measurement voids near the center of the channel (Fig. 3f), and dynamic masking yields reasonable, large velocities near the center of the channel (Fig. 3g). Note that dynamic masking was used for all particle tracking results presented in Mestre et al [15], Du et al [53], Holstein et al [90], and Hussain et al [50].

## 4 Particle tracking parameter selection

In this section, we introduce some best practices and provide general guidelines for optimizing input parameters for PTV. The goal of PTV is to obtain reliable estimates of fluid flow velocity which generally become increasingly accurate as more measurements are added. Hence, it is ideal to maximize the total number of (reliable) measurements, which can be achieved by increasing the number of particles that are tracked and/or by increasing the number of frames over which a given particle is tracked. Here we discuss three important parameters that affect the efficacy of PTV: threshold, minimum area, and maximum displacement. Figure 4a shows in blue how the mean particle track length (i.e., the average number of measurements per tracked particle) and in orange how the total number of tracks vary with each of these three parameters. Note that in all three cases, the “mean length” curve is non-monotonic with a peak, indicating that an optimal parameter value exists that maximizes the mean length of the tracks. These peaks help guide the choice of the approximate optimal parameters, which is explained in more detail in the following paragraphs. An uncertainty analysis of PTV is presented in Appendix A.

**Fig. 4.**
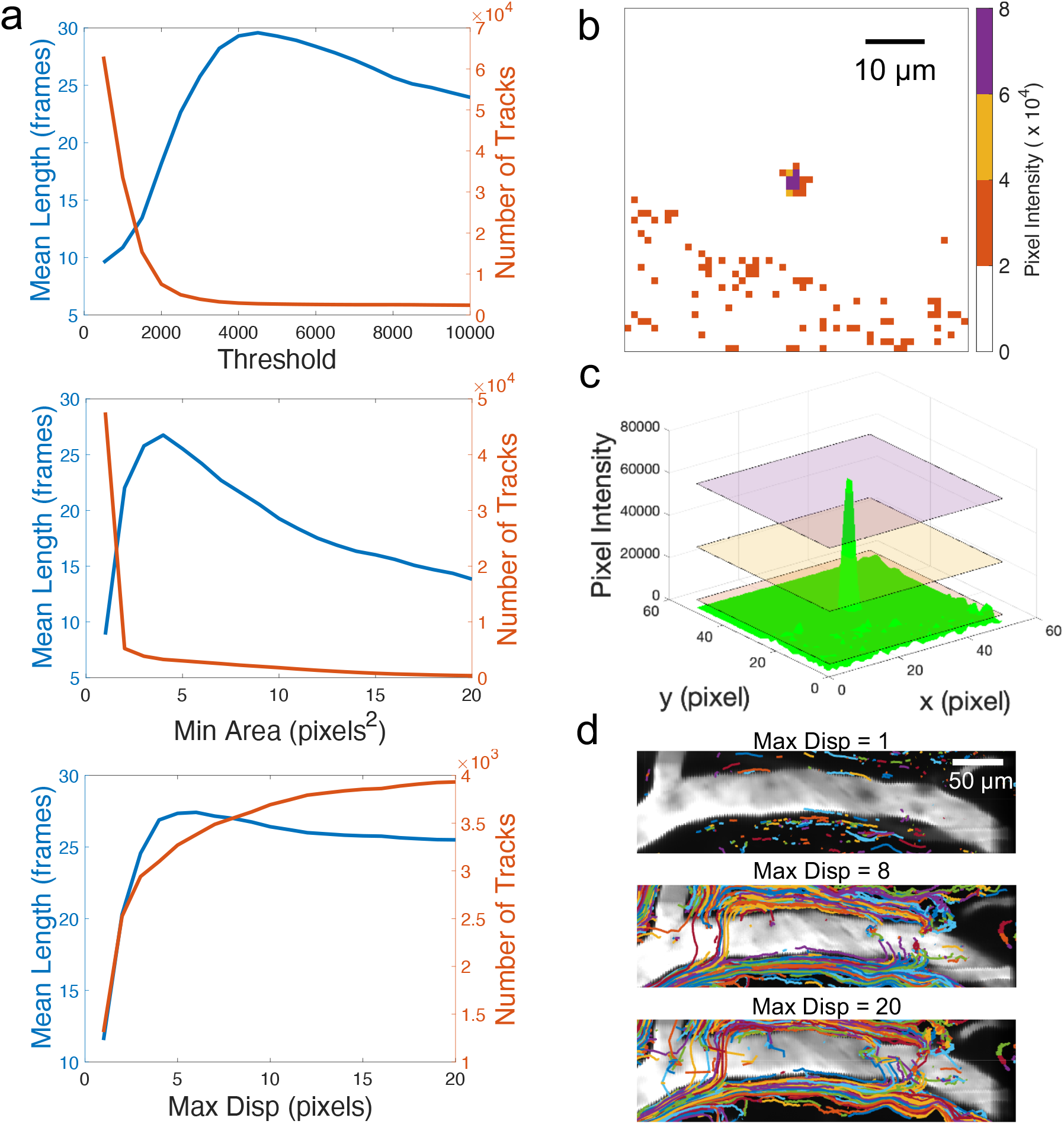
Parameter selections for PTV. **(a)** Plots with two *y*-axes characterizing the mean length of the particle tracks (blue) and the number of particle tracks (orange) for variations in (top) threshold, (middle) minimum area, and (bottom) maximum displacement. **(b-c)** Visualization of particle area as the pixel intensity threshold changes. Particles are identified more often and more reliably for lower threshold values, as long as the threshold is above the noise floor. **(d)** Higher maximum displacement increases the number of tracks, but the length of the tracks is lowered.

As mentioned above, the threshold sets a minimum pixel intensity value for identifying a particle. The two-photon microscopy images we analyze are 16-bit, meaning the pixel intensity values vary from 0 to 65535, so the threshold should also be chosen in this range. The choice of a high threshold (purple in Fig. 4b-c) will lead to smaller identified particles (since fewer pixels are above that given threshold). Conversely, the choice of a lower threshold (yellow in Fig. 4b-c) will lead to larger particles, but too low of a threshold (orange in Fig. 4b-c) will lead to erroneous identification of many particles if that threshold approaches the noise floor of the images (orange specks in the bottom of Fig. 4b; green spikes in Fig. 4c). Indeed, this effect is responsible for the large number of tracks at low threshold in Fig. 4a (top). Thus, the choice of approximate optimal parameters is more nuanced than simply maximizing the total number of measurements (equal to the mean length *×* number of tracks), which would occur for minimal threshold. By maximizing the mean track length, PTV typically achieves the greatest fidelity since the Lagrangian trajectories of particles will often pass through regions with different flow speed – long tracks suggest that certain regions are not being systematically under-sampled.

The choice of minimum particle area (“Min Area” in Fig. 4a middle) is closely tied to the threshold intensity. This parameter specifies the minimum number of contiguous pixels (adjacent and diagonally connected) that must be above the chosen intensity threshold to identify and track a given particle. Consequently, as visualized in Fig. 4b-c, smaller values of the minimum area should be chosen when higher intensity thresholds are used; similarly, larger values of minimum area can be used when the intensity threshold is low, and this choice may help reduce erroneous particle identification when the threshold begins to approach the noise floor (e.g., for a threshold of 1000 in Fig. 4b, which corresponds to orange, a minimum area of six would lead to identification of only three particles – the true particle in the center and two erroneous ones). In general, the choice of threshold and minimum area should be made together and may vary across different experiments depending on the fluorescent particle size, spatial resolution, and magnification.

The third parameter considered here is the maximum displacement (“Max Disp” in Fig. 4a bottom). As mentioned above, the particle tracking algorithm uses kinematic predictions to link and track a particle through sequential frames in time. Briefly, the maximum displacement specifies the radius over which a particle’s expected position (based on a kinematic prediction, such as the average displacement over the previous few frames) can be linked to a particle identified in the next sequential frame. Note that a kinematic prediction is not available for a newly identified particle, so the maximum displacement is applied to link particles in sequential frames based on their positions (see the “Nearest Neighbor” heuristic discussed in ref. [80]). When the choice of the value of maximum displacement is too small, new particles cannot be linked across sequential frames because their true displacement is larger than the maximum displacement, leading to very few tracks and a small mean track length (Fig. 4a bottom; also compare Fig. 4d top and middle). As the maximum displacement is increased, the number of tracks will also increase, but as the maximum displacement becomes excessively large, the rate of erroneously linked tracks will increase (compare Fig. 4d middle and bottom). When an erroneous link occurs, it has the tendency to perturb the kinematic predictions which sometimes causes the linking to fail in sequential frames. Hence, when the maximum displacement becomes excessively large, the mean track length will decrease, as seen in Fig. 4a bottom. The ideal maximum displacement varies with spatial resolution, magnification, frame rate, and fluid velocity, so it may be difficult to predict *a priori*. Ideally imaging parameters would be chosen such that the maximum displacement is large enough to not heavily rely on subpixel identification of the particle centroid (i.e., maximum displacement < 3). However, the maximum displacement should be smaller than the mean particle spacing in the image, so as to reduce the likelihood of erroneous links.

## 5 Vessel Diameter Measurements

The mechanism(s) driving CSF through PVSs in the brain are not yet well understood [34]. However, substantial evidence – both experimental [12, 24, 82, 91] and theoretical [25, 26] – suggests that arterial pulsations arising from systolic-diastolic variations in blood pressure over the cardiac cycle contribute to CSF bulk flow via pulsations in arterial diameter. In addition to systolic-diastolic variations, recent experiments [90, 92] have also demonstrated that large amplitude variations in vessel diameter occur over longer time scales due to neurovascular coupling, a process in which neuronal activity in the brain elicits arterial dilation that enhances glymphatic transport. Hence, here we describe techniques for measuring changes in vessel diameter which can be correlated with observations of changes in CSF flow characteristics, as described in refs. [12, 15, 50, 51, 53].

Figure 5a shows a TPM snapshot of a pial artery (red) with adjacent PVSs and several stagnant aggregated microspheres (green blobs); perpendicular colored lines indicate local measurements of the arterial diameter. Each measurement is obtained by analyzing the intensity profile of the fluorescence channel corresponding to the vasculature (red, in this case). We first define a line that extends beyond the boundary of the artery and is perpendicular to its axis (red dashed line in Fig. 5a). The pixel intensity along this line is then interpolated onto a one-dimensional grid with higher resolution than the raw image (typically, 10- to 100-fold), as depicted in Fig. 5b. We next identify the edges of the vessel by computing the spatial derivative of pixel intensity using a second-order accurate central finite difference (Fig. 5c). Two local maxima are then identified which we define as the edges of the vessel (red vertical lines in Fig. 5b-c). This calculation is repeated at multiple locations throughout space and in multiple frames, and can be plotted as shown in Fig. 5d. Such plots can be correlated with changes in CSF velocity, providing valuable insights into the underlying mechanism of CSF bulk flow in PVSs.

**Fig. 5.**
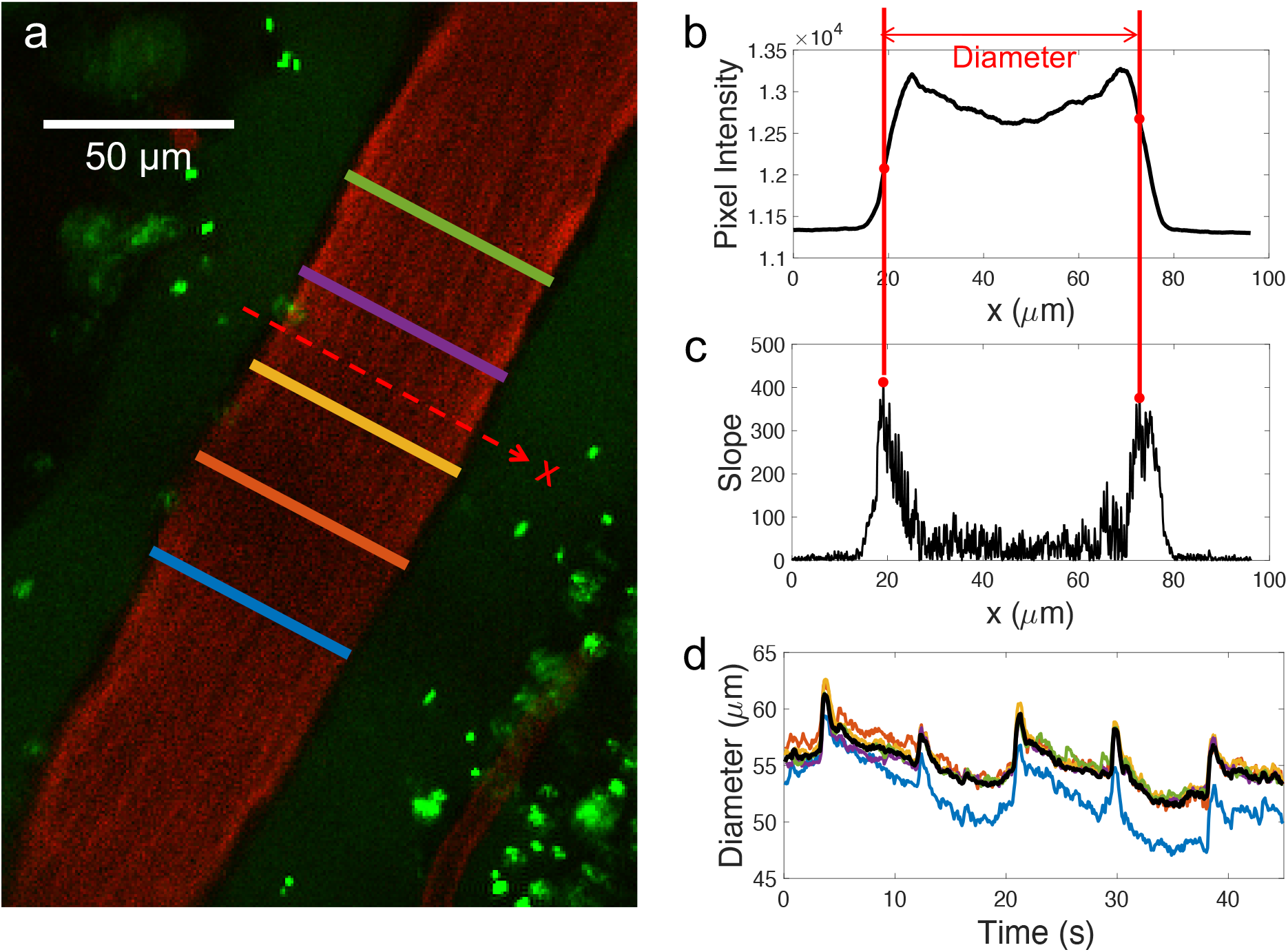
Quantitative measurements of changes in arterial diameter. **(a)** Two-photon microscopy image of a pial artery (red) and its adjacent PVS with several stagnant aggregated particles (green blobs). The solid colored lines indicate measurements of arterial diameter, while the dashed red line indicates an example region over which we interpolate the pixel intensity to identify the vessel diameter. **(b)** Plot of the interpolated pixel intensity along the red dashed line in (a). **(c)** The second-order accurate central finite difference of the pixel intensity profile in (b). The edges of the vessel are identified based on the two local maxima, indicated by the solid, vertical red lines. **(d)** Time series of the change in artery diameter with locations corresponding to the same color scheme used in panel (a). The solid black curve corresponds to the median across space evaluated at each instant of time.

## 6 Extension of Techniques to Cervical Lymphatic Vessels

The image analysis techniques previously introduced can be effectively applied to cervical lymphatic vessels, which play an important role in draining CSF from the skull [36, 37, 39, 47, 48], as discussed in the Introduction. Lymphatic vessels typically transport fluid against a mean adverse pressure difference across adjacent lymphangions (i.e., segments of lymphatic vessel separated by valves) via the combined action of contracting smooth muscle cells lining the lumen circumference and the opening/closing of bileaflet valves. This transported fluid, which includes CSF, eventually drains into the venous blood. We have recently shown that following traumatic brain injury, cervical lymphatic function becomes impaired contributing to brain edema and poor functional outcomes, but pharmacological interventions that restore lymphatic function improve functional outcomes [50]. All image analysis techniques presented above can be adapted to quantitatively measure efflux through the cervical lymphatic vessels, which we next describe.

Figure 6a shows a snapshot of a TPM recording of a cervical lymphatic vessel which has been visualized using green fluorescent dye and red fluorescent microspheres. Both tracers were injected into the CSF and within about five minutes both appear in the lymphatic vessel, demonstrating that CSF directly drains via this route. The time series of images were registered using the same method described in section 3.1, which is critically important. The neck region, which contains cervical lymphatic vessels and nodes, is in close proximity to the carotid arteries and respiratory tract which generate copious motion artifacts that must be corrected via image registration in order to obtain reliable measurements.

**Fig. 6.**
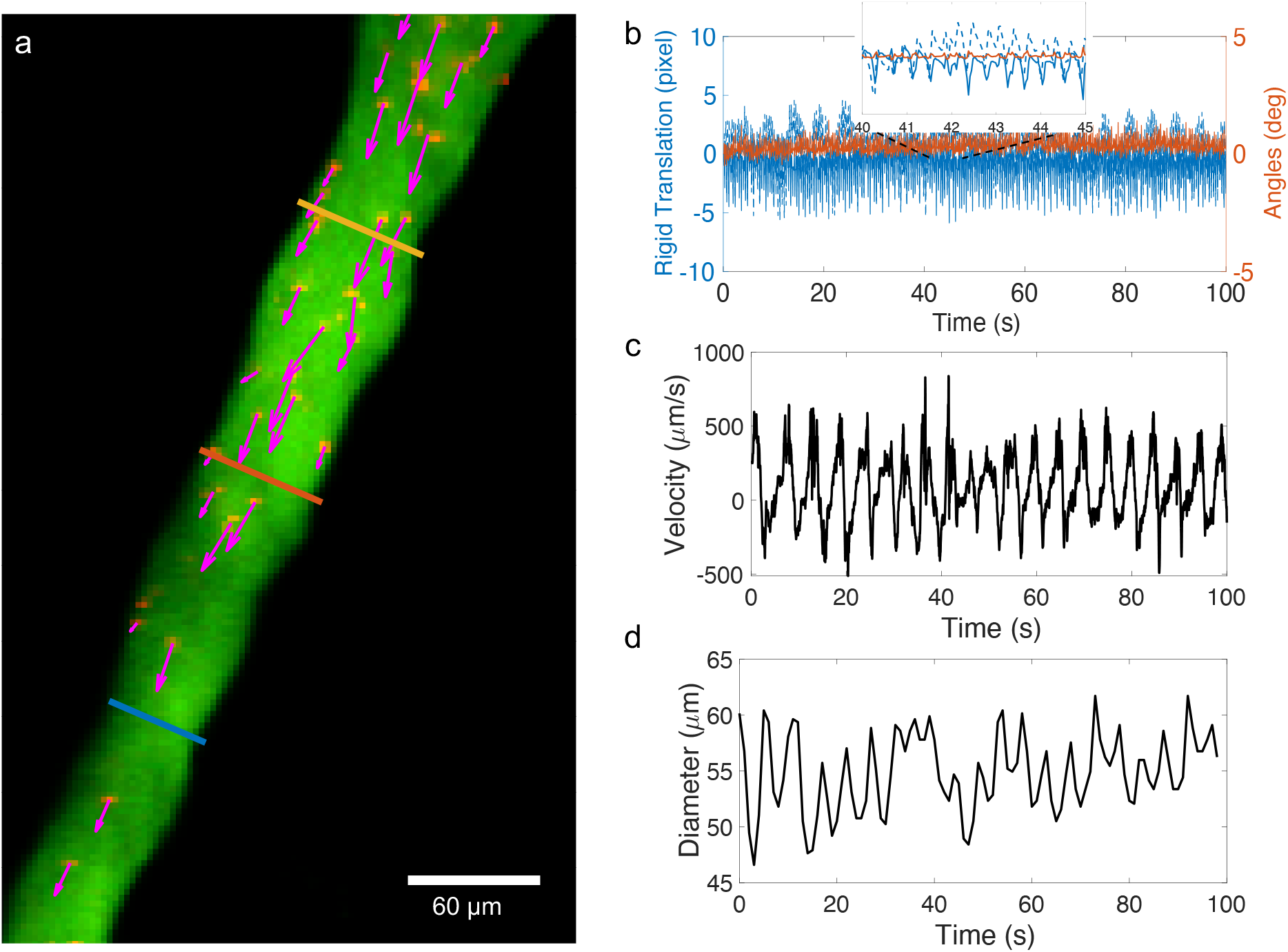
Image analysis techniques presented above, applied to a cervical lymphatic vessel. **(a)** TPM image of a cervical lymphatic vessel visualized using fluorescent dye (green) and microspheres (red). Magenta arrows represent instantaneous velocity of the particles inside the vessel. Three perpendicular colored lines indicate profile locations where the vessel diameter is measured. **(b)** A time series of registration values for rigid translation (solid and dashed blue curves) and rotation (orange curve). **(c)** A time series of the spatially-averaged instantaneous velocity obtained from particle tracking. Positive value indicate prograde flow, while negative value indicate retrograde flow. **(d)** A time series of the instantaneous vessel diameter which reveals large amplitude contractions of the vessel that closely matches the timing of changes in fluid velocity plotted in (c).

Whereas only rigid translations were adequate for measurements in PVSs (Fig. 2), we find that also including a rotational transformation greatly improves the registration quality for cervical lymphatic vessels (Fig. 6b, orange curve).

Once registration is complete, we perform particle tracking on the microspheres flowing through the cervical lymphatic vessels (red channel), enabling quantitative measurement of the spatially-averaged instantaneous flow speed (Fig. 6c). We also apply the same vessel diameter measurements introduced in section 5 to the green channel to obtain measurements of the change in cervical lymphatic vessel diameter (Fig. 6d). Such diameter measurements can be used to characterize the intrinsic pumping of lymphatic vessels [93]. Together, identifying changes in flow speed and contraction amplitude/frequency can provide crucial insights into disrupted fluid transport, as occurs following traumatic brain injury [50].

## 7 Conclusions

In this study, we outlined the image analysis techniques applied to images acquired from TPM which enables quantitative measurement of CSF flow. These scripts, as well as a short working example, are freely available online [62]. Microspheres injected into the CSF are tracked via a predictive algorithm. These particle tracks can be superimposed to estimate the size and shape of PVSs, and velocity measurements can be binned in space and time-averaged to obtain a reliable estimate of the net flow speed. However, generating high-quality PTV requires image pre-processing. To account for minor shifts in the field of view that can result in erroneous CSF flow quantification, we performed registration via rigid translation using spatial cross-correlation on a time-series of images. We highlighted a unique challenge that is encountered in this experimental protocol: some microspheres may become stuck to the boundaries of PVSs, causing erroneous zero-velocity measurements. To solve this problem, we introduced three different masking methods and reported that dynamic masking is the most promising strategy. We also offered some tips for optimizing PTV parameters including threshold, minimum area, and maximum displacement. We demonstrated that near-optimal parameter values are typically identified iteratively with the criteria of maximizing the mean track length. Furthermore, we demonstrated how changes in vessel diameter, hypothesized to be an important driving mechanism of CSF flow, can be measured in TPM images by identifying the location of the largest pixel intensity gradient to define the edges of the vessel and measure the diameter in time. Finally, we explored how the same image analysis techniques can be applied to cervical lymphatic vessels, which multiple studies indicate is an important route for CSF drainage [38, 47, 48, 50].

Our image analysis techniques for quantifying CSF dynamics offer a significant advantage in their automated tracking feature, whether for particle tracking or vessel diameter measurement. Some previous pioneering efforts have quantified CSF or lymph fluid flow through manual tracking of particles/cells [94, 95] and vessel edges across frames [95]. Such approaches are time consuming and prone to errors from operators. Our algorithm overcomes these limitations, allowing for efficient and reliable analysis of tens of thousands of frames or more, leading to accurate measurements of velocity fields and vessel diameters [12]. The image analysis techniques presented here offer various advantages over other measurement modalities, largely owing to the capabilities of TPM. In particular, TPM stands out by its ability to generate high-resolution images (e.g., 512*×*512 dimensions with 0.5 µm/pixel) while maintaining an excellent frame rate (in excess of 30 Hz) at a precise imaging depth of up to several hundred micrometers into opaque tissue. These features are valuable when monitoring the flow of microspheres within the CSF and tracking vessel wall dynamics. We highlight that application of these techniques extends beyond the brain and cervical lymphatic vessels; these tools could be utilized to quantify flow in peripheral lymphatic vessels, assess CSF circulation in the spine, analyze blood flow within the cardiovascular system, or quantify a wide range of other biological processes.

The image analysis techniques discussed here utilize 2D imaging within a precise plane, estimated to have a depth of approximately 1 µm. It is important to note that this imaging plane is typically *not* perfectly aligned with the axis of the artery, which is parallel to the primary direction of flow inside the PVS. Consequently, our measurements capture a 2D projection of a 3D flow. While 3D volumes (“z-stacks”) can be obtained using TPM by recording 2D planes at sequential depths, this approach greatly reduces the frame rate, making it unsuitable for 3D PTV applications. To achieve more precise measurements, fast 3D rastering TPM techniques are needed.

Many recent and ongoing studies aim to investigate the mechanisms by which CSF flow through PVSs is enhanced during sleep [10, 92]. The methodology presented here has previously been applied to mice that are anesthesized – not sleeping naturally [12, 15, 49, 53]. Recent studies indicate that CSF flow varies substantially across different types of anesthesia [96] and in comparison to natural sleep [97]. Training mice to fall asleep under a TPM is quite challenging but possible [92]. A promising alternative approach to more easily image CSF dynamics under natural sleep may utilize a portable miniaturized TPM [98, 99]. By attaching the portable TPM, CSF flow could be measured during normal behaviors, including natural sleep.

In this article, we have presented techniques for PTV applied in vivo in mice to quantify CSF dynamics. Such techniques have been used previously to characterize changes in CSF flow in acute arterial hypertension [12] and neurovascular coupling [90], as well as following stroke [15], cardiac arrest [51], and traumatic brain injury [50]. These direct, quantitative measurements provide both parameterization (e.g., PVS size, Reynolds number) and validation (e.g., flow speed) which can be used to improve analytical/numerical models [15, 26, 100–102]. Future measurements may help quantify flow in different compartments to help settle open questions related to CSF/ISF transport (including IPAD, glymphatic, and other hypotheses). We emphasize that the techniques outlined here enable robust and efficient in vivo measurements. Such an in vivo approach is more reliable than inferring transport rates from ex vivo analysis of fixed tissue since recent studies [12, 53, 103] have revealed anatomical changes and irregular flows generated during the tissue fixation and/or death. Measurement techniques outlined here may also prove valuable if applied to animals beyond mice, such as pigs, whose brains are gyrencephalic (i.e., have cortical folds, like the human brain) [104, 105]. Finally, in vivo PTV may help in building a mechanistic understanding of how neuromodulation enhances glymphatic flow [106–108], perhaps facilitating the development of future medical devices and pharmaceutical approaches aimed at reducing the cognitive decline associated with a wide array of neurological conditions.

## Acknowledgments

We thank Kim Boster for valuable insight and suggestions, as well as substantial contributions to the development of our MATLAB visualization tool “imagei.m”. We also thank Keelin Quirk for implementing more advanced kinematic predictions in our particle tracking MATLAB script “PredictiveTracker.m”. D.K. and J.T. are supported by a Career Award at the Scientific Interface from Burroughs Wellcome Fund. Y.G., M.N., and D.H.K. are supported by NIH BRAIN Initiative U19NS128613, NIH National Center for Complementary and Integrative Health R01AT012312, and US Army MURI W911NF1910280.

## Appendix A Particle tracking uncertainty analysis

In this Appendix, we analytically calculate the approximate uncertainty in particle tracking velocity measurements using propagation of error based on estimated uncertainties in particle position and image acquisition time. The numbers used here are specific for one particular data set [62], serving as a concrete example, but the reader could readily repeat these calculations for different imaging parameters. This derivation applies to two-photon microscopy, wherein each 2D image is assembled over a finite window in time during which each pixel is individually rastered. Hence, for images acquired at 29.6 Hz, each acquired image has an uncertainty of approximately Δ*t* = 33.8 ms. Note this value is an upper bound for the temporal uncertainty, this upper bound will generally be larger/smaller when a greater/lesser number of pixels are recorded, and more precise (smaller) values of Δ*t* could be obtained by accounting for details of the TPM rastering technique.

The velocity of a tracked particle in frame *n* can be estimated, to first order, as 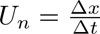, where Δ*x* is the measured displacement of the particle since frame *n−* 1 and Δ*t* is the measured time elapsed since frame *n −* 1. Denoting the error in Δ*x* as *u_x_* and the error in Δ*t* as *u_t_*, the error in *U_n_* is

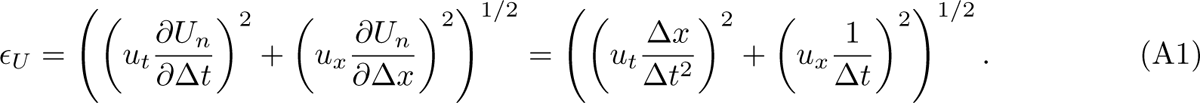

Our image processing algorithm locates each particle by finding the centroid of a contiguous bright region above a given threshold, typically achieving error on the order of 0.1 pixel. In this data, each pixel has lateral dimension 1.04 µm, so we estimate *u_x_ ≈* 0.104 µm. As mentioned above, for a frame rate of 29.6 Hz, Δ*t* = 33.8 ms. Now suppose the root-mean-square single-frame displacement is Δ*x* = 2.158 µm (this value is obtained empirically by performing particle tracking). Estimating the timing error *u_t_* for a two-photon microscope is subtle because images are produced not by making simultaneous measurements from an array of sensors but by making subsequent measurements as the single focal point rasters the field of view. If the focal point traces column-by-column, then measuring a particle moving to an adjacent column involves greater timing error than measuring a particle moving to an adjacent row. Considering image dimensions 512 *×* 512 pixels, the two timing errors would be roughly Δ*t/*512 and Δ*t/*512^2^, respectively. To be conservative, we take *u_t_ ≈* Δ*t/*512 = 66.0 µs. Using these values, we find *ɛ_U_* = 3.08 µm/s.

In practice, we estimate the velocity with a higher-order method, convolving the measured position with a kernel that provides differentiation and smoothing. Explicitly, the numerical scheme used in our code makes the estimate

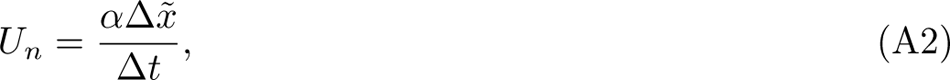

where *α* = 2*/*(*π*^1*/*2^erf(3) *−* 6*e^−^*^9^) *≈* 1.129 and

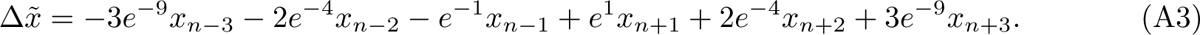

The velocity error is

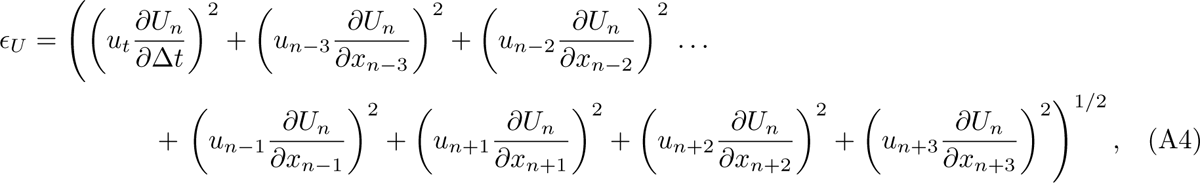

where *u_n−_*_3_ is the measurement error associated with location *x_n−_*_3_, *u_n−_*_2_ is the measurement error associated with location *x_n−_*_2_, and so on. Assuming homogeneity implies that all those errors have the same value, which we again denote *u_x_*. Then, the velocity error becomes

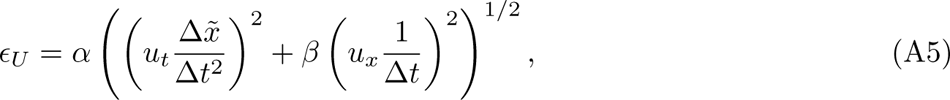

where *β* = 18*e^−^*^18^ + 8*e^−^*^8^ + 2*e^−^*^2^ *≈* 0.273. To estimate the value of Δ*x*°, we consider the case in which a particle’s displacement between any two frames is the measured root-mean-square value Δ*x*, implying *x_n_*_+1_ *− x_n−_*_1_ = 2Δ*x*, *x_n_*_+2_ *− x_n−_*_2_ = 4Δ*x*, and *x_n_*_+3_ *− x_n−_*_3_ = 6Δ*x*. Therefore, Δ*x*° = *γ*^1*/*2^Δ*x,* where *γ* = (2*e^−^*^1^ + 8*e^−^*^4^ + 18*e^−^*^9^)^2^ *≈* 0.782. Altogether, the velocity error is

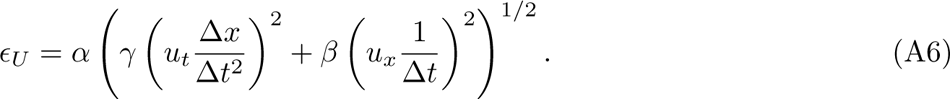

Comparing to Eq. (A1), we see that the error in velocity estimated with the higher-order numerical scheme differs from the error in first-order velocity estimates only by factors of order unity. Again taking the same values for *u_t_*, *u_x_*, Δ*t*, and Δ*x*, the velocity error in the higher-order scheme is *ɛ_U_* = 1.82 µm/s, about 40% lower than with the first-order estimate.

